# DNetPRO: A network approach for low-dimensional signatures from high-throughput data

**DOI:** 10.1101/773622

**Authors:** Nico Curti, Enrico Giampieri, Giuseppe Levi, Gastone Castellani, Daniel Remondini

## Abstract

The objective of many high-throughput “omics” studies is to obtain a relatively low-dimensional set of observables - signature - for sample classification purposes (diagnosis, prognosis, stratification). We propose DNetPRO, *Discriminant Analysis with Network PROcessing*, a supervised signature identification method based on a bottom-up combinatorial approach that exploits the discriminant power of all variable pairs. The algorithm is easily scalable allowing efficient computing even for high number of observables (10^4^ − 10^5^). We show applications on real high-throughput genomic datasets in which our method outperforms existing results, or compares to them but with a smaller number of selected variables. Moreover the linearity of DNetPRO allows a clearer interpretation of the obtained signatures in comparison to non linear classification models

---

The huge amount of high-dimensional omics data (e.g. transcriptomics through microarray or NGS, epigenomics, SNP profiling, proteomics and metabolomics, but also metagenomics of gut microbiota) poses enormous challenges as how to extract useful information from them. One of the prominent problems is to extract low-dimensional sets of variables - signatures - for classification and diagnostic purposes, for example to better stratify patients for personalized intervention strategies based on their molecular profile [9, 2, 6, 1].

Many approaches are used for these classification purposes [3], such as Elastic Net [5], Support Vector Machine, K-nearest Neighbor, Neural networks and Random Forest [8]. Some methods select signature variables by means of single-variable scoring methods [7, 4] (e.g. inferential testing for two-class comparison), while others search for projections in variable space, and then perform a dimensionality reduction by thresholding the projection weights, but these approaches could fail even in simple two-dimensional situations (Supplementary Fig. 1a).

Our method - DNetPRO, *Discriminant Analysis with Network PROcessing* - generates multivariate signatures starting from all variable pairs tested with Discriminant Analysis (scheme of the algorithm pipeline in Fig. 1). The computing time for variable space exploration is proportional to the square of the number of variables (ranging from 10^3^ to 10^5^ in a typical high-throughput omics study), but the method provides an alternative approach to single-feature selection methods, and provides a hard-thresholding approach at difference with projection-based variable selection methods. Moreover, the geometrical simplicity of the resulting class-separation surfaces allows an easier interpretation of the results, as compared with very powerful but black-box methods like nonlinear-kernel SVM or Neural Networks. This linear separation might not be common in some classification problems (e.g. image classification) but it is very plausible in biological systems, in which many responses to perturbation consist in increase or decrease of variable values (e.g. expression of genes or proteins, see Supplementary Fig. 1b).

**Figure 1:**
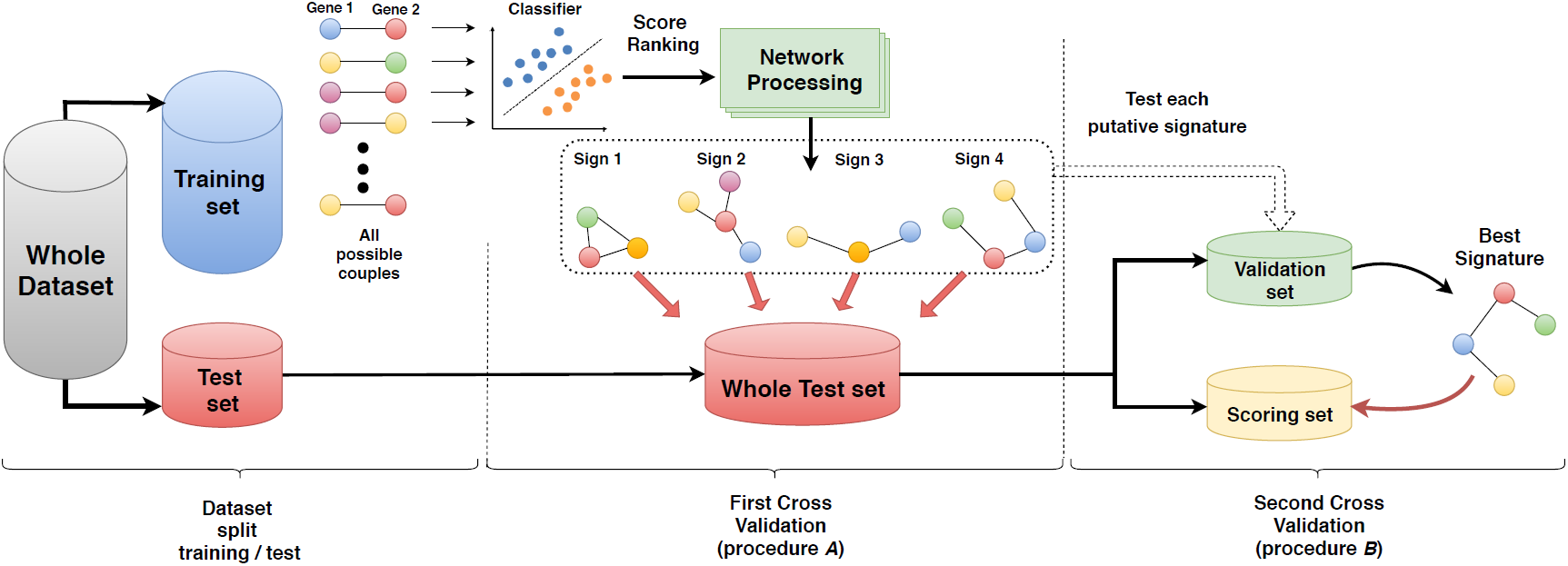
Scheme of DNetPRO algorithm. On the “training set;, all possible pairs of variables are used for Discriminant Analysis, generating the fully connected network with links weighted by pair classification performance. By thresholding the weighted links, several signatures come out as connected components, and then their performance is evaluated on the “whole test set; (procedure *A*). A unique best signature can be identified on a “validation set; and tested in a “scoring set;, obtained by further splitting the “whole test set; (procedure *B*).

We then applied our method to real biological data, core sets extracted from the The Cancer Genome Atlas (accession number syn300013, doi:10.7303/syn300013), used in a previous study [10] which aimed at quantifying the role of different omics data types (e.g. mRNA and miRNA microarray data, protein levels measured with Reverse Phase Protein Array - RPPA) via different state-of-the-art classification methods. This allowed us to compare our results to a large set of commonly used classification methods, by using their performance validation pipeline (accession number syn1710282, doi:10.7303/syn1710282). For each cancer type (*kidney renal clear cell carcinoma KIRC, glioblastoma multiforme GBM, ovarian serous cystadenocarcinoma OV and lung squamous cell carcinoma* LUSC) we show results on mRNA data (see *Supplementary Material* for the results on miRNA and RPPA datasets).

We remark that DNetPRO can provide more than one signature as a final outcome, given by all the connected components found in the variable network, or a unique top-performing signature can be obtained by a further cross-validation step (procedure *A* and procedure *B* in Fig. 1, respectively).

The results are shown, as distribution of AUC (Area under the curve) score, in Fig. 2 (a) for the best signatures obtained with procedure *A* (corresponding to the validation approach used in [10]), while results with the full cross-validation procedure *B* are shown in Fig. 2 (b). As expected, performances decrease with the introduction of the second cross validation step, but the values remain quite stable showing the robustness of the extracted signatures, and we remark that the validation procedure used in the reference paper by Yuan et al. resembles our approach without the second validation step.

**Figure 2:**
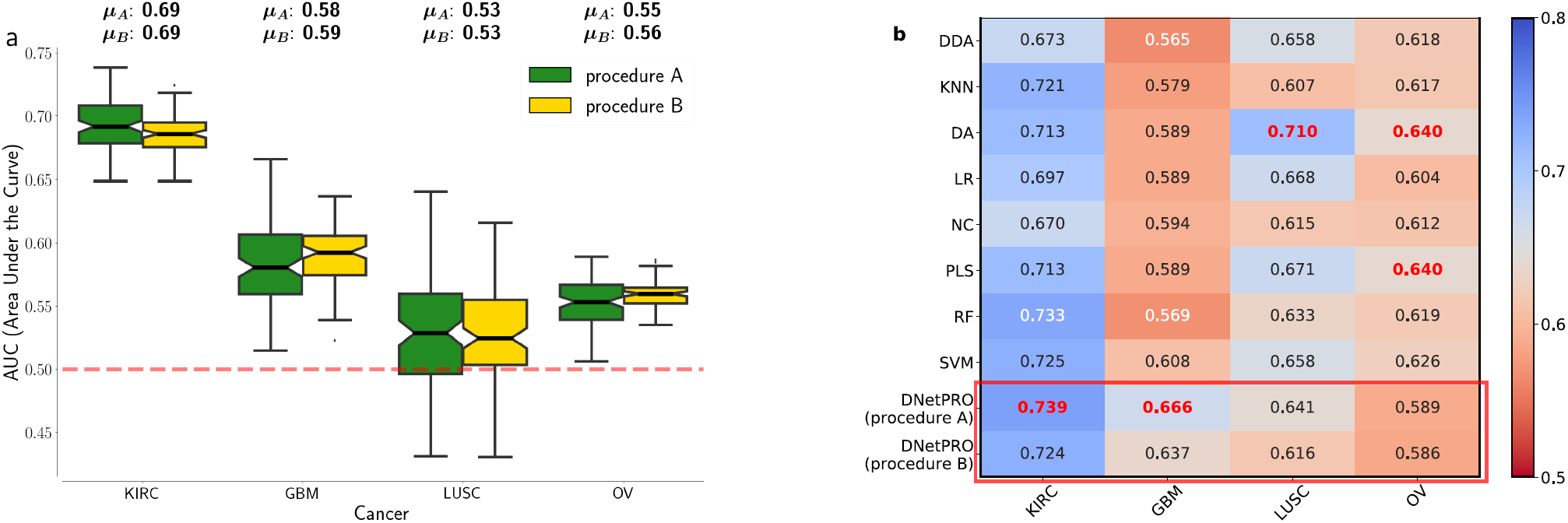
Results obtained by the DNetPRO algorithm pipeline on four mRNA tumor datasets, as from the Synapse database[10]. (a) Distributions of AUC values for the tumor datasets. Green boxplots: results using procedure *A* as described in Fig. 1; yellow boxplots: results obtained using procedure *B*. (b) Comparison of DNetPRO results with the methods used in the paper of Yuan et al.: max AUC values obtained over the 10-Fold cross-validation procedure.

All results are comparable (LUSC) or better (KIRC, GBM) than the results reported in [10], except for the OV dataset, also with the more conservative approach involving a further cross-validation step. The size of the extracted signatures is quite constant, and smaller than 500 genes in each pipeline execution. Analogous results are obtained also for the miRNA dataset, in which our method outperforms in three over four cases, while the RPPA dataset shows less significant results (Supplementary material).

To test the robustness of our method, since each cross-validation procedure may generate different signatures, we measured the overlap of the genes belonging to each mRNA signature over 100 simulations with different training-test data splitting.

The method we presented has several advantages: easy scalability on parallel architectures, simple signature interpretation allowing a valuable application in a biomedical context and a significant robustness in a highly noisy environment such as genomics measurements.

## METHODS

*Any information about the datasets used and the implementation of the algorithm is available in the on-line version of the paper.*

## ACKNOWLEDGEMENTS

The authors acknowledge IMI-2 HARMONY n. 116026 EU Project and IMforFUTURE Horizon 2020 (EU) Project.

