## Supplementary Materials for "DNetPRO: A network approach for low-dimensional signatures from high-throughput data"

[1]Department of Physics and Astronomy, University of Bologna
[2]Department of Experimental, Diagnostic and Specialty Medicine, University of Bologna
[3]INFN Bologna

### DNetPRO algorithm description

Given a *dataset*, consisting in $S$ *samples* (e.g. cells, patients) with $N$ observations each (our *variables*, e.g. genes or protein expression profiles) the signature identification procedure is summarized with the following pipeline:

- separation of available data into a *training* and a *test* set (typically 33/66, or 20/80);
- estimation of the classification performance on the training set of all $S(S-1)/2$ *variable pairs* through a computationally fast and reproducible cross-validation procedure (leave-one-out cross validation was chosen);
- selection of top-performing pairs through a hard-thresholding procedure. The performance of each pair constitutes a *weighted link* of a network in which nodes are the variables connected at least through one link;
- every *connected component* in which the network is divided into constitutes an identified classification signature.
- (optional) in order to reduce the size of an identified signature, the *pendant nodes* of the network (i.e. nodes with degree equal to one) can be removed, in a single step or recursively until the core network (i.e. a network with all nodes with at least two links) is reached.
- all signatures are applied onto the test set to estimate their performance (this is the procedure $A$ in Fig. 1 of the main text).
- to identify the best performing signature, a further cross validation step can be performed, with a further dataset splitting into training set (to identify best pairs and possibly multiple signatures), test set (to identify the best signature) and validation set (to evaluate best signature performance). This is the procedure $B$ in Fig. 1 of the main text.

To test the performance of all variable pairs, we used a diag-quadratic Discriminant Analysis, a robust classifier that allows fast computation. We remark that a variant of DNetPRO method can be used also for dimensionality reduction of network structures, as can be obtained by correlation-based approaches [7, 4].

### Algorithm implementation

In DNetPRO algorithm, every analysis of a variable pair is an independent computational process, so it is easily parallelizable to increase speed performance. We thus developed a multi-core parallelized version of the algorithm, with shared memory architecture and with the analysis process divided into many threads.

To improve the scalability of our algorithm, we implement the pipeline scheme using Snakemake [5] rules (Python codes) and the computation of discriminant networks (the most time expensive step) is performed in C++ with the OpenMP multi-threading support. An example of the pipeline work-flow for a single cross-validation step is shown in Fig. 1.

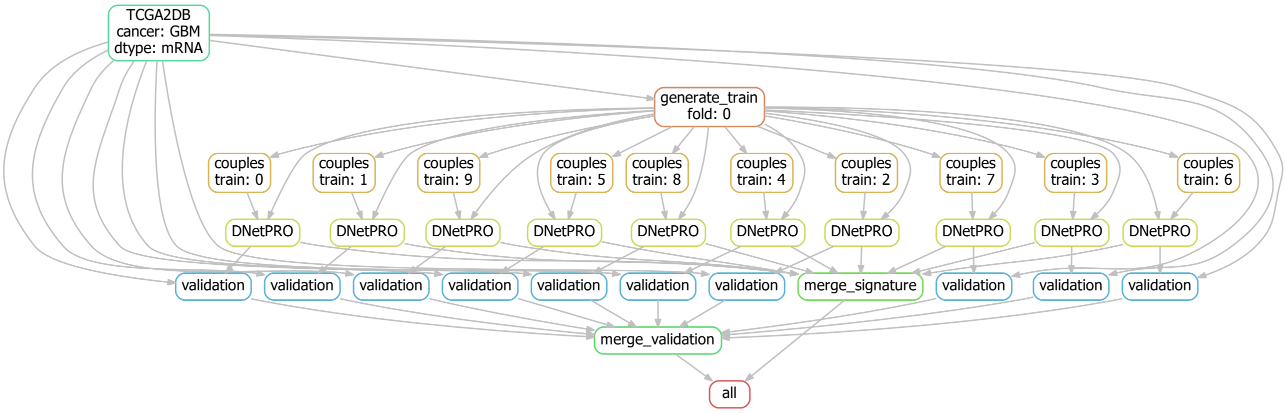


Figure 1: Example of DNetPRO pipeline with a single cross validation step. It is highlighted the independence of each fold from each other. This scheme shows a possible distribution of the jobs on a multi-threading architecture or for a distributed computing architecture. The second case allows further parallelization scheme (hidden in the graph) for each internal step (e.g. the evaluation of each pair of genes).

With Snakemake support, we can easily distribute our pipeline on several machines using a simple master-slave scheme. In this case each step of Fig. 1 can be performed by a different computer unit preserving the multi-threading steps, with a maximum scalability and the possibility to enlarge the problem size and the number of variables.

We tested our algorithm implementation on a dedicated HPC server (128 GB RAM memory and 2 CPU E5-2620, with 8 cores each) and on a common laptop (8 GB RAM memory and 1 CPU i7-6500U, with 2 cores) using the same dataset (20530 probes and 150 samples, with a total of more than $2 \times 10^8$ combination of variables). Both tests were done with the hyper-threading enabled and considering either the evaluation of the full combination of variables either the ordering of them. The pair evaluation takes around 1 minute on the first architecture and 10 minutes on the second one, showing a good scalability of the code on the available threads. We have to remark that the reading of the dataset is done in serial mode and the sorting algorithms are drastically affected by the cache size, giving a huge advantage to the server machine. We also provided an MPI (*Message Passing Interface*) version of the code to allow analyses of large datasets with a greater number of variables but it was not necessary to use it in this context since the multi-threading version on a bioinformatics server was fast enough for our applications.

The combinatorial algorithm developed is also wrapped in a Python class using Cython [2] to allow a good integration also with other common tools of machine learning and the most common used python libraries like scikit-learn library [8]. We also tested our Cython implementation against a parallel Python implementation, monitoring its scalability across different dataset sizes and number of available threads (ref. Fig. 2). The provided DNetPRO implementation is more than $10^4$ faster than the Python one with the same dataset size and thus it can be used also with higher number of variables.

The pipeline implementation and the entire set of codes developed in this work are available at DNetPRO-pipeline).

### DnetPRO and data characteristics

Methods that select variables for multi-dimensional signatures based on single-variable performance can have limits in predicting higher-dimensional signature performance. An example is shown in Fig. 3(a), in which both variables taken singularly perform poorly, but their performance becomes optimal in a 2-dimensional combination, through a simple linear separation of the two classes.

It is known that complex separation surfaces characterize classification tasks associated to image and speech recognition, for which Deep Networks are successfully applied in recent times, but in many cases biological data, such as gene or protein expression, are more likely characterized by a up/down-regulation behavior (as shown in Fig. 3(b) top), while more complex behaviors (e.g. a "windowed" optimal range of activity, Fig. 3(b) bottom) are much less common. Thus, discriminant-based methods (and logistic regression methods alike) can very likely provide good classification performances in these cases (as demonstrated by our results with DNetPRO) if applied in at least two-dimensional spaces. Moreover, the "linearity" of these methods (that generate very simple

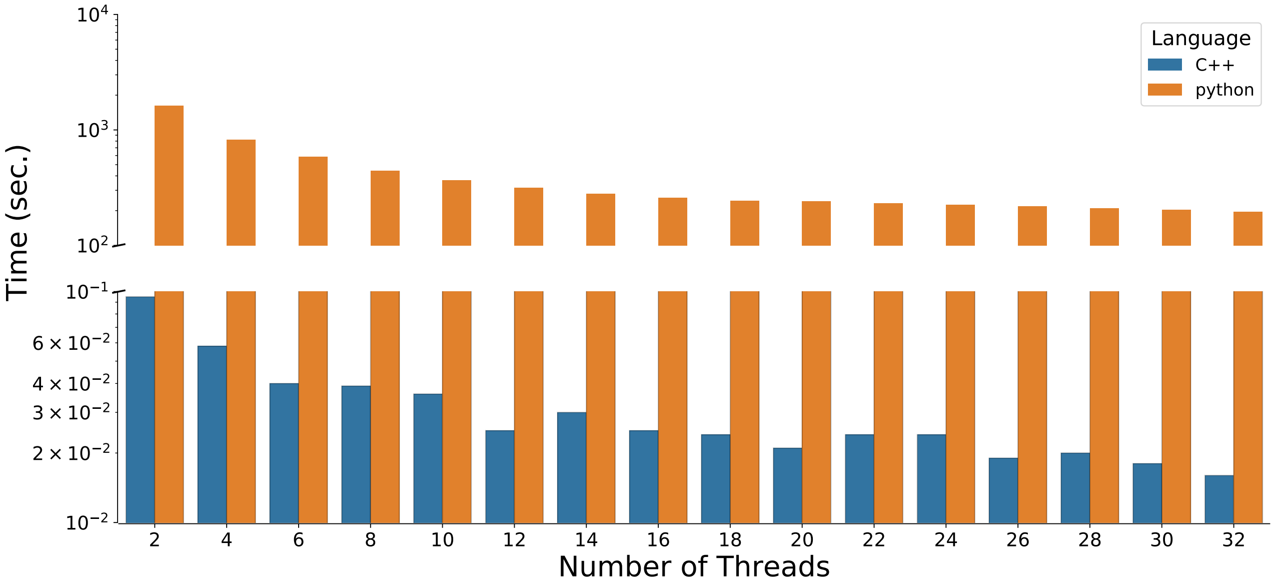


Figure 2: Scalability of DNetPRO algorithm implementation across various number of threads, using a dataset composed by 90 samples and 90 variables. We compare the computational time performances of our Cython version of the DNetPRO algorithm against a parallel pure-Python one. The tests were performed on a HPC server (128 GB RAM memory and 2 CPU E5-2620, with 8 cores each). The Cython implementation is more than $10^4$ faster than the pure-Python implementation. The trend oscillations of Cython times are due to an unbalanced partition of the tasks related to the number of samples provided.

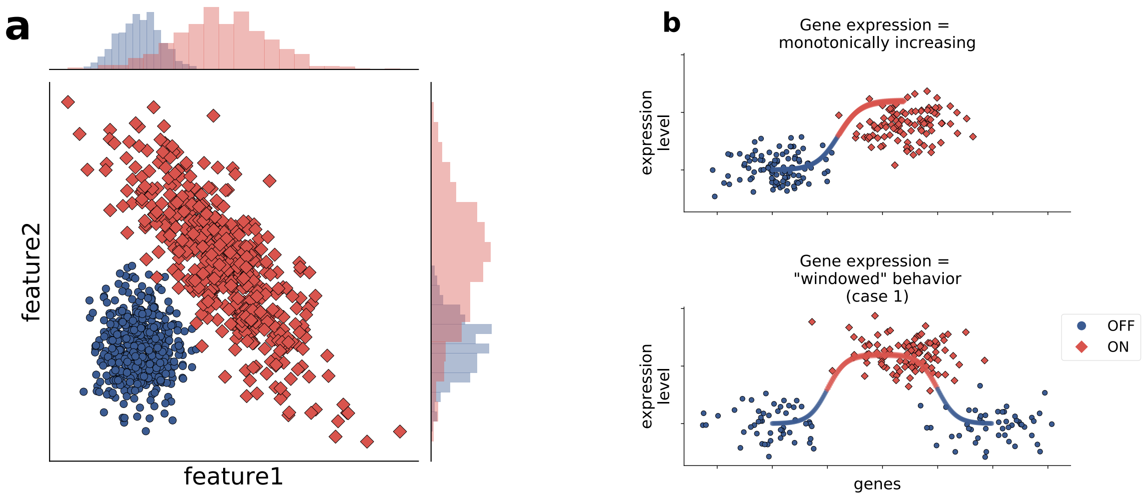


Figure 3: (**a**) An example in which single-variable classification performance fails in predicting higher-dimension classification performance. Both variables (*feature1* and *feature2*) badly classify in 1-D, but have a very good performance in 2D. Moreover, classification can be easily interpreted in terms of relative higher/lower expression of both probes. (**b**) Activity of a biological feature (e.g. a gene) as a function of its expression level: top) monotonically increasing, often also discretized to an on/off state; center, bottom) "windowed" behavior, in which there are two or more activity states that don't depend monotonically on expression level. X axis: expression level, Y axis, biological state (arbitrary scales).

class separation surfaces, i.e. linear or quadratic) guarantee that a "buildup" of a multidimensional signature based on 2-dimensional initial signatures (our variable pairs) is feasible.

We remark that the discriminating signatures have a purely statistical relevance, being generated with a purpose of maximal classification performance, but previous applications of a simplified version of the Dnet-PRO algorithm [1, 9, 3, 10] allowed to gain knowledge on the biological mechanisms associated to the studied phenomenon.

### Ranked pairs analysis and signature characterization

Since the number of variable pairs is typically very large (e.g. $10^8$ pairs with gene expression microarrays containing about $10^4$ probes) many of them may achieve the same performance, since the possible performance values are integer number typically in a limited range (corresponding to the number of available samples, $10^2 - 10^3$ in many cases). Therefore, the ranking of pairs by their performance is characterized by multiple "plateaus"(Fig. 4(a)), and the selection of probe pairs based on a hard thresholding procedure is highly influenced by this feature.

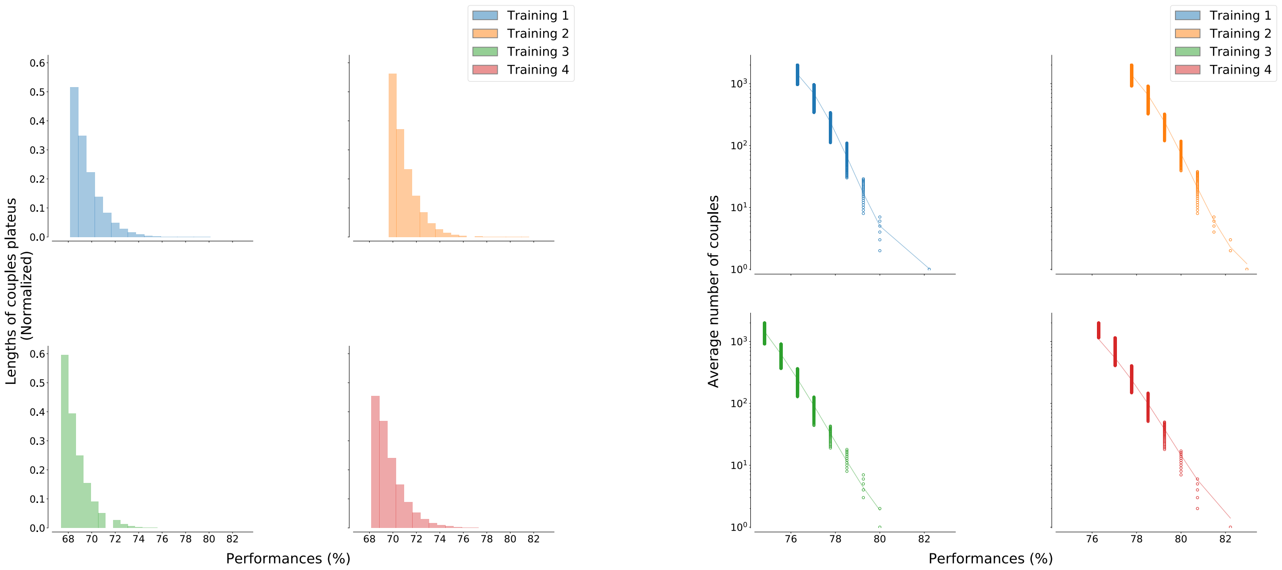


Figure 4: Analysis of ranked pairs distributions according to the performance score obtained in the training step. (**a**) The distribution of plateau lengths is approximately exponential. (**b**) Average number of pairs with the same score value: this behavior is typical in ranking order distribution and it can be fitted by the relation $f(x) = A(M + 1 - r)^b/r^a$ as shown in [6], where $r$ is the rank value, $M$ its maximum value, $A$ a normalization constant and $(a, b)$ two fitting exponents.

As in other cases of ranked values [6], we can fit these ranking distributions with a combination of power-law functions obtaining a good agreement with experimental points (Fig. 4(b)).

We also observed that "star"-networks frequently appear, with one variable highly connected to many others which are only connected with it. This happens when a variable has a strong discriminating power, to which other possibly less relevant variables get linked for noisy fluctuations.

As stated before, we suggest that these variables (pendant nodes in the star network) can be removed from the signature without significantly affecting its performance. The procedure can be applied for one single step (in order to remove nodes pending from a star configuration) or it can be applied recursively, until the signature becomes constituted only by the 2-core network (i.e. with all nodes having degree $\geq 2$). Empirical analysis performed on real data has shown that the removal of these variables does not affect significantly the signature performance, and in the meanwhile it allows a significant reduction of its dimensionality. Since there is no clear theoretical explanation of this behavior, we suggest to introduce this step only optionally, since it is not easy to quantify the risk of loosing relevant information from the removed variables.

The underlying idea is that the more connected are the nodes, the more the variables in the signature "work well" together, a plausible hypothesis given the linear sample separation surface provided by the Discriminant classifier. Moreover, the network structure of the signature suggests further considerations about the relevance of a variable as a function of its role in the network (e.g. node centrality such as degree or betweenness centrality).

### Description and processing of the Synapse dataset

The TCGA (The Cancer Genome Atlas) core sets of data used are available at the Synapse homepage (accession number syn300013, doi:10.7303/syn300013).) created by Yuan et al. [11], and are composed of four tumor datasets: kidney renal clear cell carcinoma (KIRC), glioblastoma multiforme (GBM), ovarian serous cystadenocarcinoma (OV) and lung squamous cell carcinoma (LUSC).

The summary description of the datasets used is reported in Table 1.

| Cancer | mRNA | miRNA | Protein | Number of samples |
|---|---|---|---|---|
| GBM | AgilentG4502A | H-miRNA_8x15k | RPPA | |
| | 17814 | 533 | [a] | 210 |
| KIRC | HiseV2 | GA+Hiseq | RPPA | |
| | 20530 | 1045 | 166 | 243 |
| OV | AgilentG4502A | H-miRNA_8x15k | RPPA | |
| | 17814 | 798 | 165 | 379 |
| LUSC | HiseqV2 | GA+Hiseq | RPPA | |
| | 20530 | 1045 | 174 | 121 |

Table 1: Description of the benchmark dataset used in the paper. The type of cancer, the platform with the number of probes and the number of samples per tumor type are shown.
[a] Missing data.

Each tumor dataset was pre-processed by adding a zero-mean Gaussian random noise ($\sigma = 10^{-4}$)) to remove the possible null values in the database, which could produce numerical errors in the evaluation of the distance between gene/protein expression values. Then, we randomly split each dataset in training and test sets with a stratified (i.e. balanced for class sample ratio) 10-fold procedure: with the stratification we are reasonably sure that each training-set is a good representative of the whole sample set. The choice of a 10-fold splitting is aimed to reproduce the analysis pipeline presented by Yuan et al. [11] with an analogous cross-validation procedure. Since we don't have exact details of their data splitting, the cross validation was repeated 100 times, for a total of 1000 training procedures for each tumor (OV, LUSC, KIRC, GB) and data type (mRNA, miRNA, RPPA). Each training procedure led to the extraction of multiple signatures.

We chose threshold values in order to obtain a resulting number of variables in the signatures in the order of $10^2 - 10^3$, and identified all connected components of the network as signatures. If more than one component existed, each one was considered as a different signature.

The final multidimensional signatures were tested by a Discriminant Analysis with a diag-quadratic distance, to avoid possible problems about covariance matrix inversion (as for the Mahalanobis distance in which there is a higher number of coefficients to be estimated from the data).

In the single cross-validation pipeline (procedure $A$ in Fig. 1 of the *paper*) the best signature was extracted as the one reaching the highest accuracy score during the training step.This best signature was then tested over the available test set.

When also the second cross-validation step was included in the pipeline (procedure $B$ in Fig. 1 of the *paper*) the best signature was chosen as the best performing over a subset of the whole test set (renamed *test set*), and the final performance was evaluated on the remaining *validation set*.

To compare our results with the work of Yuan et al. [11], we used the AUC (*Area Under the Curve*) score, that they provided in the paper as the result of their analyses. The distribution of our results could be compared to the single score value given in their work.

### Analysis of miRNA and RPPA datasets

The same analysis pipeline presented in the paper about gene expression data from the Synapse dataset is applied in this Section also to the miRNA and RPPA datasets, with the results presented in Fig. 5.

The results obtained on the miRNA dataset (Fig. 5 (a, b)) are comparable to the reference, while for the RPPA dataset only the LUSC tumor shows AUC values comparable with the others. Moving from the procedure $A$ to the procedure $B$, i.e. adding a second cross-validation step, the RPPA performances drastically decrease for the KIRC and OV, while their remain quite stable for the LUSC dataset. The same behavior is shown in the miRNA datasets in which however both performances are still comparable or better (KIRC, GBM, LUSC) than the reference ones.

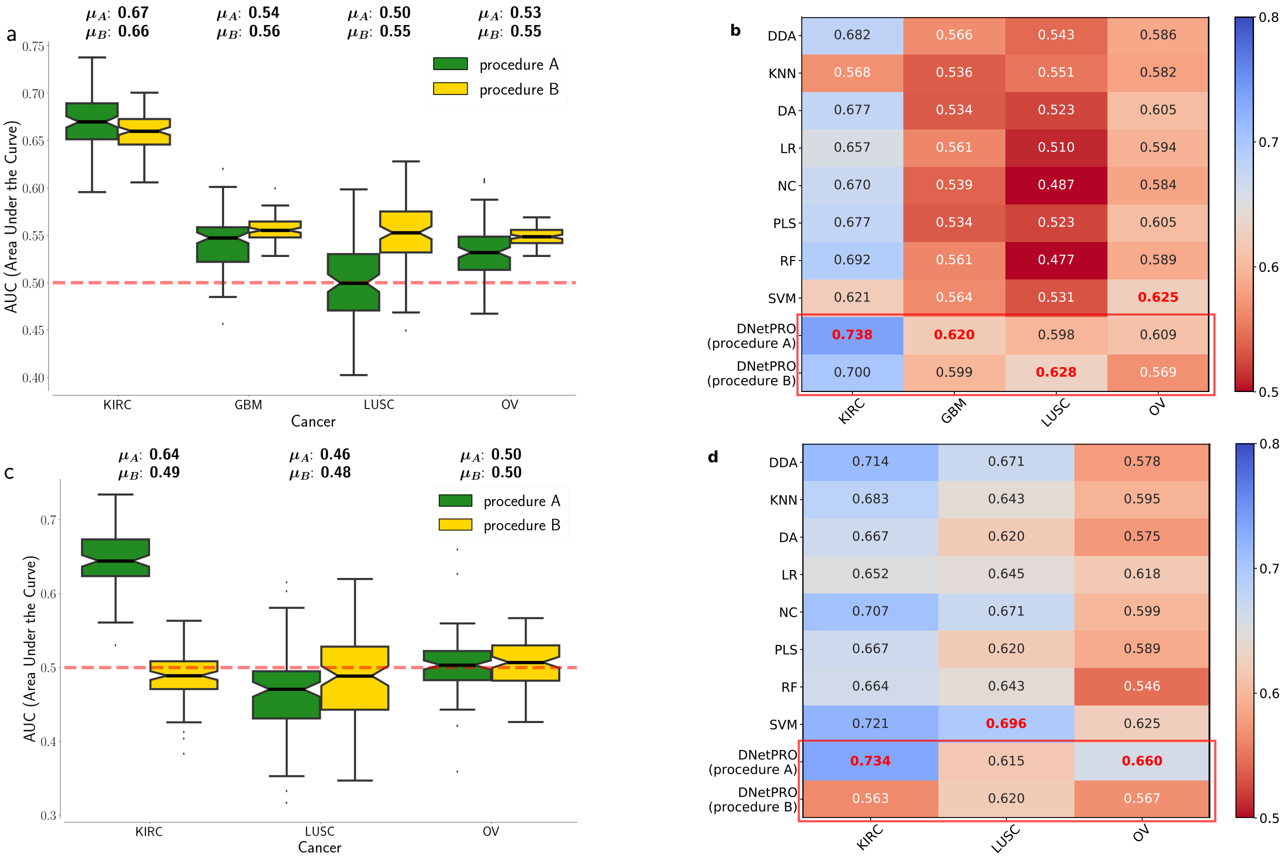


Figure 5: Results obtained by the DNetPRO algorithm pipeline on the four Synapse miRNA and RPPA tumors datasets. **(a, c)** Distributions of AUC scores obtained over the four datasets. Green boxplots: results using procedure $A$ of DnetPRO; yellow boxplots: results obtained using procedure $B$. **(b, d)** Comparison of DNetPRO with the methods used in [11]. The reported values are the max AUC values obtained over the 10-Fold cross-validation procedure.
